# Climate adaptation shaped by subtle to moderate allele frequency shifts in loblolly pine

**DOI:** 10.1101/635862

**Authors:** Amanda R. De La Torre, David B Neale

**Affiliations:** School of Forestry, Northern Arizona University, AZ86011, USA; Department of Plant Sciences, University of California-Davis, One Shields Avenue CA95616, California, USA

**Keywords:** GEA, loblolly pine, local adaptation, high-density linkage map

## Abstract

Understanding the genomic basis of local adaptation is crucial to determine the potential of long-lived woody species to withstand changes in their natural environment. In the past, efforts to dissect the genomic architecture in gymnosperms species have been limited due to the absence of reference genomes. Recently, the genomes of some commercially important conifers, such as loblolly pine, have become available, allowing whole-genome studies of these species. In this study, we test for associations between 87k SNPs, obtained from whole-genome re-sequencing of loblolly pine individuals, and 270 environmental variables and combinations of them. We determine the geographic location of significant alleles and identify their genomic location using our newly constructed ultra-dense 26k SNP linkage map. We found that water availability is the main climatic variable shaping local adaptation of the species, and found 492 SNPs showing significant associations with climatic variables or combinations of them. Our results suggest that adaptation to climate in the species might have occurred by many changes in the allele frequency of alleles with moderate to small effect sizes, and by the smaller contribution of large effect alleles in genes related to moisture deficit, temperature and precipitation. Genomic regions of low recombination and high population differentiation harbored SNPs associated with principal components but not with individual climatic variables, suggesting climate adaptation might have evolved as a result of different selection pressures acting on groups of genes associated with an aspect of climate rather than on individual climatic variables.

## INTRODUCTION

Local adaptation may arise by differential selection pressures across heterogeneous environments leading to increased fitness in the local environment compared to the non-local environment. Although of great interest by population geneticists, the genomic architecture of local adaptation remains largely unsolved in natural populations of non-model species (Anderson et al., 2013). A majority of studies aiming to dissect the genetic architecture of local adaptation have focused at detecting signals of selection in which a new advantageous mutation in a single gene is rapidly driven to fixation, also known as the “hard sweep” model (Smith & Haigh, 1974). In contrast, recent genome-wide association studies in humans and forest trees species have suggested a largely polygenic basis of local adaptation (Hancock et al., 2010; Pritchard et al., 2010a; Neale & Kremer, 2011; Le Corre & Kremer, 2012). If a population that is well adapted to a geographic location moves to a new environment, natural selection will increase the frequency of certain alleles until the typical phenotype in the population matches the phenotype optimum in the new environment (Pritchard et al., 2010a). This type of adaptation also called “polygenic adaptation” is characterized by subtle to moderate shifts in allele frequencies and may be frequent in traits that have standing genetic variation for selection to act on, are highly heritable and controlled by many loci of small effect (Pritchard et al., 2010a, 2010b; Berg & Coop, 2014).

A common approach to find genes contributing to local adaptation has been based on the idea that genes under selection should be more genetically differentiated among populations than a neutral locus, and will therefore have high Fst values (Cavalli-Sforza, 1966; Whitlock & Lotterhos, 2015). Loci mostly affected by spatially heterogeneous selection will have high Fst values, whereas the ones under spatially uniform balancing selection will show lower than neutral Fst values (Whitlock & Lotterhos, 2015). Due to a wide distribution of Fst values in neutral markers, only advantageous alleles with high frequencies can be detected under the Fst outlier approach (Pritchard et al., 2010). This problem is exacerbated by the lack of power of individual tests when correcting for multiple comparisons over thousands of loci. As a result, weakly selected loci, characteristic of polygenic adaptation, will unlikely to be detected (Le Corre & Kremer, 2012; Yeaman, 2015; Whitlock & Lotterhos, 2015). In contrast, genotype by environment association (GEA) studies are more likely to detect signals associated with smaller allele frequency shifts (Hancock et al., 2010; Forester et al., 2018).

Lately, increasing numbers of genome-wide markers have enabled the study of the genomic architecture of local adaptation in natural populations. The geographic and genomic distribution of adaptive alleles can give us insights into the evolutionary forces that had shaped adaptation in a species. For example, alleles associated with day length in natural populations of *Arabidopsis thaliana* showed narrow geographic distribution in which one allele had rapidly driven to high frequency in the population as a result of a hard-selective sweep, whereas SNPs associated with relative humidity had widespread distributions (Hancock et al., 2011). Also, the clustering of adaptive alleles in genomic regions of low recombination due to linkage or divergence hitchhiking can give us insights into the maintenance of population differentiation and local adaptation in the face of gene flow (Via, 2012; Yeaman, 2013).

In the past, efforts to dissect the genomic architecture of local adaptation in gymnosperms species have been limited due to the absence of reference genomes. Recently, the genomes of some commercially important conifers have become available, allowing whole-genome studies of these species (De La Torre et al., 2014a; Neale et al., 2017). Current assemblies of reference genomes are still quite fragmented and do not allow the location of genes in the genomes unless high-density linkage maps are available. Loblolly pine (*Pinus taeda*) is a widely distributed species in the southeastern United States, characterized by its outcrossing mating system, large population sizes, weak population structure and rapid decay of linkage disequilibrium (Eckert et al., 2010b). Phylogeographic studies of unglaciated North America suggested loblolly pine follows the Mississippi River discontinuity, which is consistent with a dual Pleistocene refugial model, and has been used to explain differences in growth, disease resistance, drought tolerance and genetic differentiation between eastern and western populations (Soltis et al., 2006; Eckert et al., 2010a, 2010b; Teskey et al., 1987; Schmidtling, 2001). In this study, we aimed to dissect the genomic architecture of climate adaptation in loblolly pine. We tested for associations between 270 environmental variables and 87k SNPs obtained from widely distributed coding and non-coding regions across the 22-Gb genome of the species. We then determined the geographic and genomic location of significant alleles using our newly constructed, ultra-dense, 26k SNP linkage map. In addition to identify the main climatic variables driving the adaptation of the species, we were also interested in the following questions: a) Are adaptive alleles globally occurring alleles with varying frequencies or localized ones? b) Do adaptive alleles have narrow or widespread genomic distributions? c) Does local adaptation occur by large or subtle shifts in allele frequencies?.

## MATERIAL AND METHODS

### Sample collection and SNP genotyping

Needle tissue from 377 outcrossing, unrelated individuals distributed across the species’ natural range were collected from the ADEPT2 common garden located in Mississippi, southeast United States (Figure 1A). In addition, two three-generation full-cross outbred pedigrees constructed and maintained by the Weyerhauser Company were used to collect 192 needle samples (Sewell et al. 1999). From these, 92 full-sib progeny samples came from the *qtl* pedigree, and 100 full-sibs came from the *base* pedigree. DNA was extracted using a protocol that included one day of tissue lysis and incubation at 96°C, followed by several steps of precipitation and filtering using the Qiagen DNeasy mini-prep Plant kit with an Eppendorf automated pipetting workstation. DNA concentration and quality were evaluated using picogreen on a Qubit Fluorometer. Raw reads from whole-genome re-sequencing data for ten individuals were used to call a large number of SNPs (455M) that were later scored, filtered and included in an Affymetrix Axiom myDesign species-specific and customized SNP array comprising 635k SNP markers (full description of this procedure can be found in De La Torre et al., 2018). After removing SNPs that did not pass the genotyping quality control criteria and those that were monomorphic, we kept 84,738 high-resolution SNPs. In addition, 3,087 gene-based SNPs previously reported in Eckert et al. 2010 were added to the dataset, resulting in a total of 87,825 SNPs. From these SNPs, 20,367 matched genes, exons, transcripts or a combination of those.

**Figure 1.**
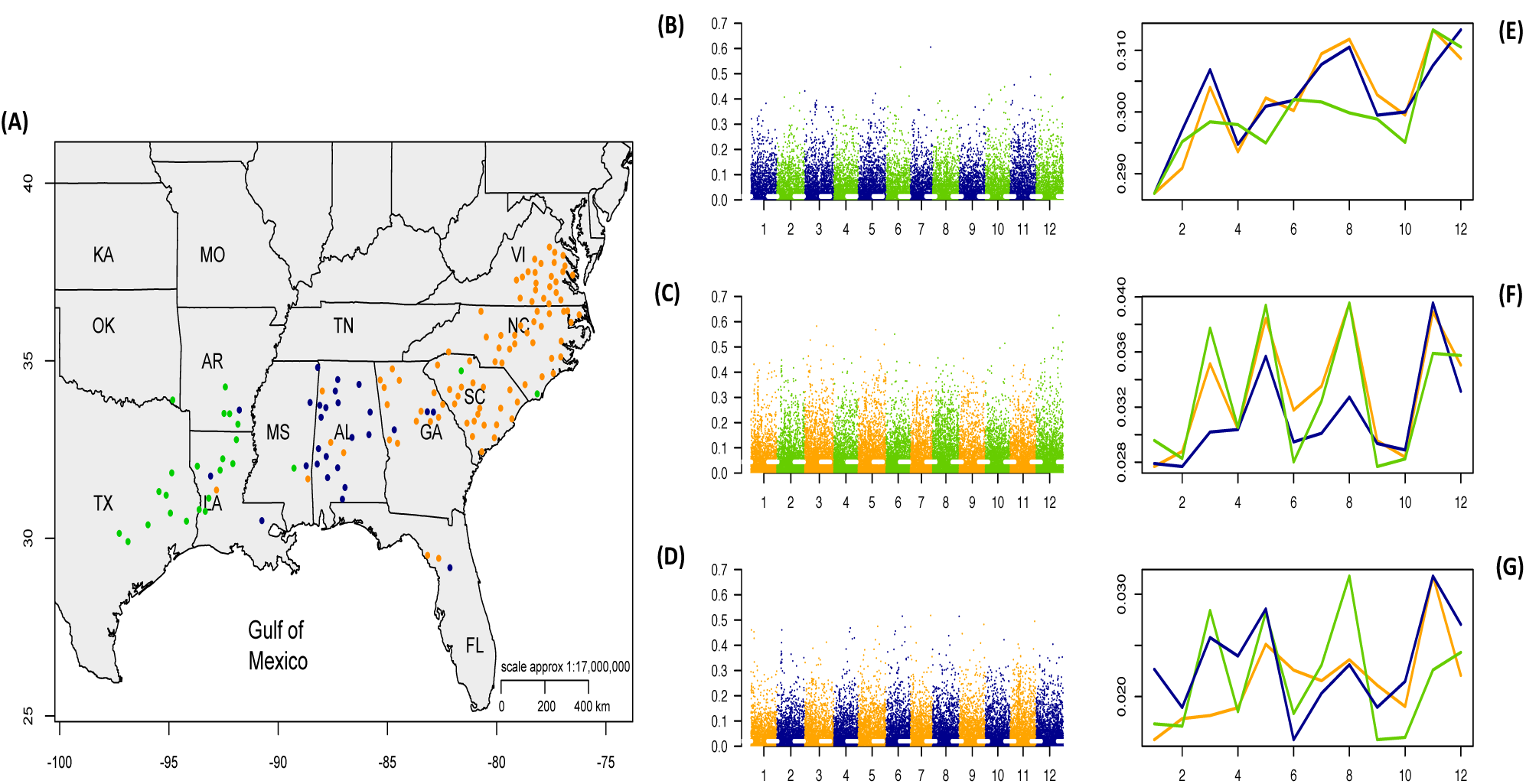
Population structure and nucleotide diversity of loblolly pine based on 87k genome­wide SNP markers. Results of PCA and Structure suggest the presence of three different clusters along longitude (Fig.1A). Manhattan plots show pairwise Fst distributions between west and center populations (Fig. 1B), west and east populations (Fig.1C), and center and east populations (Fig. 1D). Dotted horizontal white line represents the mean pairwise Fst for each pair of populations. Average nucleotide diversity, Fst one vs all populations, and pairwise Fst among populations are shown in Figures 1E, 1F, 1G. Colors in all figures match genetic clusters (populations) in Fig.1A.

### Population structure, diversity estimates and Fst outlier test

Within populations’ nucleotide diversity for each SNP in each of the 12 linkage groups was estimated with the R Package PopGenome v.2.2.4 (Pfeifer et al., 2014). Population pairwise fixation index (Fst) was estimated by comparing all possible pairs of populations, and by comparing each population against all other individuals (Fst one vs. all). Fst values for each SNP in each linkage group were plotted in Figures 1B, 1C and 1D; and average Fst values were displayed in Figures 1E, 1F, 1G. Population structure in the SNP dataset was evaluated using the Python2.x fastStructure algortithm based on a variational framework for posterior inference of K clusters (Raj et al., 2014). Models in fastStructure were replicated 10 times with K from 1 to 10 using the default prior; seeds for random number generators were modified for each run. The chooseK.py python script in fastStructure was used to estimate the model complexity that maximizes marginal likelihood and the model components were used to explain structure in the data. Main pipeline and Distruct package in CLUMPAK (Kopelman et al., 2015) were used for the summation and graphical representation of FastStructure results. In addition, we did a PCA analysis using the Adegenet v2.0.1 R package (Jombart, 2008; Jombart & Ahmed, 2011). Outlier SNPs showing higher Fst than neutral loci were identified using the OutFLANK R package. OutFLANK infers the null distribution by removing loci in the top and bottom 5% of the distribution, and it is suggested to be robust to demographic history (Whitlock & Lotterhos, 2015).

### Genotype-environment associations (GEA)

We used a combination of 248 monthly, seasonal, and annual variables obtained from climate normal data from 1961-1990 in ClimateNA v5.41 (Wang et al., 2016). In addition, we used 19 GIS-derived bioclimatic variables from WorldClim 2.5-min (www.worldclim.org), and aridity index by quarter (every 4 months starting January), as previously calculated by Eckert et al. (2010b). Geographical variables for each individual tree (latitude and longitude), and combinations of environmental data in the form of principal components, were also added to the analysis. Genotype by environment associations were identified for each of the 270 climatic variables with 87,859 SNPs with compressed mixed linear model (Zhang et al., 2010) implemented in the GAPIT R package (Lipka et al., 2012). To reduce the chance of identifying false positive associations as a result of population structure, we conducted the association analysis with only those SNPs having a minor allele frequency (maf) higher than 10% and used principal components of genetic data as co-variants. Manhattan plots were built using the SNP locations in our newly constructed ultra­dense linkage map for loblolly pine with the R package qqman (Turner et al., 2014). A subset of environmental traits showing significant associations with at least ten SNPs, were used to test associations between effect size (proportion of the variance explained) and minor allele frequency. SNP functional annotations were obtained from the annotated genome of loblolly pine v2.01 in TreeGenes (https://treegenesdb.org/Drupal). For the SNPs matching transcripts, we aligned them against the non-redundant protein sequences database using BLASTX 2.8.0 (evalue <1e-10) (Zheng et al., 2000).

### Latent Factor Mixed Models (LFMM) analysis

In addition, we used a Bayesian bootstrap approach implemented in LFMM command line version 1.5 (Frichot et al., 2013). LFMM accounts for random effects due to population structure and spatial autocorrelation with the use of latent factors (k) (Frichot et al., 2013). Each run was repeated five times with random seeds using the following parameters: k=3, 10,000 iterations, 5,000 burning length. Correction for multiple testing was done by adjusting p-values with the genomic inflation factor (l= median(Z-score^2^)/0.456) after combining z-scores obtained from multiple runs with the LEA R package (Frichot & Francois, 2015). Runs were repeated with k=2 and k=5, to check for sensitivity of results to the number of latent factors in the model. To address the potential correlations among environmental variables, we run a PCA analysis using the prcomp function in R package. The first 3 Principal Components resulting from this analysis were tested for associations with all the 87k SNPs in LFMM. Only SNPs with a minor allele frequency >10% were included. We also tested for the presence of genomic clusters (more than 10 SNPs within a 1cM window) for all SNPs significantly associated with any climatic variable or principal component of them. Top 20% SNPs with the highest −log10(P) values for each principal component were retained for further analysis and considered as candidates for adaptation, following Yoder et al. 2014.

### Environmental co-association network analysis

In addition to the univariate GEA analysis implemented in LFMM and Gapit, we also tested for the multivariate response of groups of genes associated with the environment, using the environmental co-association network analysis as described in Lotterhos et al. 2018. SNPs showing univariate associations in the GEA Gapit analysis were used to construct a network analysis using a hierarchical clustering of the associations between SNP allele frequencies and environmental variables using the reshape2 and gplots R packages in RStudio (RStudio Team, 2016).

### Construction of individual pedigree linkage maps

*Qtl* and *base* pedigrees’ pseudo-testcrosses were used as R/qtl (Broman et al., 2003) objects to construct ultra-dense linkage maps using the MSTmap algorithm implemented in the ASMap v.0.4 R package with default p-value (Taylor & Butler, 2017). Pairwise recombination (r) and LOD scores (obtained from linkage disequilibrium test) were estimated for each pair of SNP markers. Markers showing an r<0.5 were considered located in the same linkage group. SNP markers with pairwise recombination frequency estimated as zero (co-locating markers), were placed into a recombination bin with ASMap. Several rounds of stringent filtering included the removal of markers with 30% or more missing data, duplicated individuals, double crossovers, markers with distorted segregation patterns, and markers not mapping well in any of the linkage groups. The presence of switched alleles (wrong phase) was also accounted for. JointMap v5.0 (Van Ooijen et al., 2006) was used for fine mapping and ordering of bins obtained from ASMap.

### Construction of average-sex and consensus maps

To allow the construction of averaged-sex maps for each pedigree, a set of anchor markers composed by 131 fragment-based markers (RAPD, RFLPs, ESTs, and SSRs) and 2804 SNPs (Martinez-Garcia et al., 2013; Eckert et al., 2010b) were added to our dataset. SNPs with suspect linkages as defined by a recombination frequency higher than 0.6 and a LOD score higher than 1 were excluded in further analyses. Forty-eight individual maps were merged to create twenty-four averaged-sex maps for each pedigree. Averaged-sex maps were merged using maximum intervals (K) from 1 to 8, generating eight consensus maps for each of the 12 linkage groups, with the R package LPmerge (Endelman & Plomion, 2014). Consensus maps with the lowest root mean-squared error (RMSE) standard deviation (mapping conflicts between individuals maps and consensus) were selected for each linkage group (Endelman & Plomion, 2014). Graphical display was done with Circos (Krzywinski et al., 2009). All markers were anchored to the reference genome of loblolly pine v2.01 (https://treegenesdb.org/Drupal) with the BWA-MEM algorithm in BWA (http://bio-bwa.sourceforge.net). Convergence and map accuracy was evaluated by comparing genome physical location (scaffold ID) and linkage group; and by comparing our maps with previously published maps in the species (Martinez-Garcia et al., 2013; Westbrook et al., 2015).

## RESULTS

### Population structure and genetic diversity levels

Results of the PCA analysis with 87k SNPs implemented in the Adegenet and Gapit R packages suggest the presence of three major genetic clusters (east, center, west) that extend longitudinally across the species’ natural range (Figure 1A, Supporting Information Figure S1). When using the posterior inference of clusters based on variational Bayesian framework implemented in fastStructure, we found two major clusters when K varies from 2 to 4. When K=5, there is a third smaller cluster (Supporting Information Figure S2). However, our fastStructure test of the model complexity that maximizes marginal likelihood suggested K=2 better explains the genetic structure of the species. When K=2, the center and eastern populations are differentiated from the western populations, suggesting the Mississippi river as the major barrier for gene flow, as previously observed in De La Torre et al. 2018. We found that eastern and western populations present higher pairwise Fst estimates and were therefore more genetically distant than eastern and center, and center and western populations (p-value<0.001; Table 1; Figures 1B, 1C and 1D). For example, from the 622 SNPs with a pairwise Fst >0.3, 394 SNPs differentiated the western and eastern populations. Pairwise Fst between pairs of populations was found to significantly vary in all linkage groups with the exception of LG 7 and 10 (p-value<0.001; Table 1; Figures 1F, 1G). Nucleotide diversity across all linkage groups was significantly different among populations (p-value<0.01), and also differed between the center and west, and the east and west in linkage groups 3, 4, 7, 9 and 12 (p-value<0.001; Table 1; Figure 1E). Average nucleotide diversity for all populations, based on SNP data, was 0.296, average Fst values (one population vs all others) was 0.029, and average pairwise Fst among all pairs of populations was 0.027 (Table 1).

**Table 1.** Population differentiation (Fst comparison between one vs. all populations, and pairwise Fst calculations) and nucleotide diversity based on SNPs with known linkage group location in loblolly pine. Population labels (pop 1-3) represent genetic clusters from east to west, as described in Figure 1.

| LG | Nucleotide diversity |  |  |  | Fst one vs all |  |  |  | Pairwise Fst |  |  |  |
| --- | --- | --- | --- | --- | --- | --- | --- | --- | --- | --- | --- | --- |
|  | pop1 | pop2 | pop3 | All | pop1 | pop2 | pop3 | All | pop1/<br>pop2 | pop1/<br>pop3 | pop2/<br>pop3 | All |
| 1 | 0.287 | 0.281 | 0.273 | 0.280 | 0.028 | 0.018 | 0.029 | 0.025 | 0.016 | 0.040 | 0.015 | 0.023 |
| 2 | 0.291 | 0.292 | 0.283 | 0.289 | 0.029 | 0.018 | 0.027 | 0.025 | 0.018 | 0.039 | 0.012 | 0.023 |
| 3 | 0.304 | 0.304 | 0.287 | 0.298 | 0.035 | 0.020 | 0.036 | 0.031 | 0.018 | 0.051 | 0.017 | 0.029 |
| 4 | 0.294 | 0.290 | 0.287 | 0.290 | 0.030 | 0.020 | 0.029 | 0.027 | 0.019 | 0.041 | 0.015 | 0.025 |
| 5 | 0.303 | 0.297 | 0.283 | 0.294 | 0.038 | 0.026 | 0.038 | 0.034 | 0.025 | 0.051 | 0.018 | 0.031 |
| 6 | 0.300 | 0.298 | 0.292 | 0.297 | 0.032 | 0.019 | 0.027 | 0.026 | 0.023 | 0.041 | 0.010 | 0.024 |
| 7 | 0.310 | 0.305 | 0.292 | 0.302 | 0.033 | 0.020 | 0.031 | 0.028 | 0.021 | 0.045 | 0.013 | 0.026 |
| 8 | 0.312 | 0.308 | 0.289 | 0.303 | 0.040 | 0.023 | 0.038 | 0.033 | 0.024 | 0.055 | 0.015 | 0.031 |
| 9 | 0.303 | 0.295 | 0.288 | 0.295 | 0.030 | 0.019 | 0.027 | 0.025 | 0.021 | 0.038 | 0.012 | 0.024 |
| 10 | 0.300 | 0.296 | 0.283 | 0.293 | 0.028 | 0.019 | 0.027 | 0.025 | 0.019 | 0.038 | 0.014 | 0.024 |
| 11 | 0.313 | 0.304 | 0.306 | 0.308 | 0.039 | 0.029 | 0.034 | 0.034 | 0.032 | 0.045 | 0.021 | 0.032 |
| 12 | 0.309 | 0.311 | 0.303 | 0.307 | 0.035 | 0.023 | 0.034 | 0.031 | 0.022 | 0.047 | 0.017 | 0.029 |
| All | 0.302 | 0.299 | 0.289 | 0.296 | 0.033 | 0.021 | 0.032 | 0.029 | 0.021 | 0.044 | 0.015 | 0.027 |

### Fst outlier analysis

We screened out all SNP loci having low heterozygosity due to their confounding effects on Fst distribution, and we ended up with a dataset of 68k SNPs. Our results identified 152 SNPs with p-values less than 0.001, however none of them passed the threshold after correction for multiple testing (q value <0.01). When increasing the RightTrimFraction (parameter that removes the loci mostly affected by selection before estimating the shape of the Fst distribution through likelihood), we found that 392 SNPs passed the q value threshold, however the fit of the null distribution model was reduced. OutFLANK identified a large number of SNPs with very high Fst values of up to 0.6, however the wide distribution of Fst values did not allow a clear distinction between putatively neutral and putatively under selection SNPs, as it is expected in a Fst outlier distribution.

### Genotype-environment associations (GEA) and environmental co-association network analysis

A total of 2432 significant associations were found among 193 climate variables and 630 SNPs (FDR-Adjusted p-values<0.01). After filtering SNPs with a minor allele frequency lower than 10%, 1279 associations were found among 186 climate variables and 113 SNPs (Supporting Information Tables S1 and S2). From the 113 SNPs, 31 SNPs came from coding regions, and 50 were located in linkage groups. No significant associations between maf and effect size [0.1-0.6, mean=0.36] was found in the dataset, suggesting a low contribution of rare alleles (Supporting Information Table S2). In fact, SNPs associated with principal components of climatic variables had an average maf of 0.365, and the ones associated with climate variables had an average maf of 0.278.

From the climate variables, CMD_wt (winter Hargreaves Climatic Moisture Deficit, which is the difference between a reference evaporation and precipitation during winter) and CMD02 (Hargreaves Climatic Moisture Deficit in February) showed the highest number of associated markers, with 72 SNPs associated to each of them. All other climatic variables had a range of 3 to 13 SNPs associated to each climatic variable. Only a small group of SNPs matching transcripts aligned to known proteins at the NCBI non-redundant protein sequences database. Associated SNPs were mainly found to be involved in transport, transcriptional activity and enzymatic functions (Supporting Information Table S2). SNPs associated with any of the climatic variables were widely distributed across all 12 linkage groups in the genome of loblolly pine and some of them were associated with several climatic variables.

Hierarchical clustering of significant associations between SNP allele frequencies and environmental variables suggests the presence of two main modules, one related to aridity (temperature variables, radiation and degree-days above 18°C) and another mainly associated with humidity and cold (climate moisture deficit during winter CMDwt, NFFD, Annual Heat-Moisture index AHM) (Supporting Information Figure S4). In the first module, six SNPs (AX-172791235, AX-173011888, AX-172909447, AX-173250534, AX-173348038, and AX-173361850) showed strong associations with a large number of temperature-related environmental variables (Supporting Information Table S2). These SNPs were also found to be associated with PC1 in both the Gapit and LFMM analysis. Interestingly, even though we were not able to map any of these SNPs in our linkage map, we know that three of them are located in the same scaffold (super 3645) based on the latest genome assembly of the species. The second module is composed by several sub-modules or sub-groups. In one of them, all 72 SNPs associated with CMDwt and CMD02 cluster together. Smaller sub-modules are associated with NFFD during april (9SNPs), Eref during the summer (4 SNPs); and Annual heat moisture index (4SNPs). Although our results suggest pleiotropic effects of the some of the SNPs, our conclusions are limited by the confounding effects of environmental correlations (Supporting Information Figure S5).

PCA analysis of environmental variables showed PC1 explains 66.56% of the variation in the dataset, PC2 explains 14.49%, and PC3 6.51%. PC1 negatively correlates with Temperature-derived variables, Relative Humidity (RH) and Radiation (Rad); and positively with Degree-days below 0c (DD_0), Degree-days below 18c (DD_18), Precipitation as snow (PAS), Frost-Free period (FFP) and Temperature seasonality variables such as Isothermality (BIO3); Temperature seasonality (BIO4); Annual Temperature Range (BIO7), and Continentality (TD). PC2 positively correlates with Relative Humidity (RH), and negatively with Temperature seasonality variables (TS) (Figure 2A, Supporting Information Figure S3). LFMM results showed 716 SNPs associated with PC1, 444 SNPs with PC2, and 741 SNPs with PC3 (Bonferroni-Holmes-Adjusted p-value<0.01, maf>0.1). Changing the number of latent factors (k) that account for population structure mainly identified the same group of associated SNPs (data not shown). We selected the top 20% significant SNPs with highest −log_10_(p-value) for each PC as potential candidates for local adaptation, resulting in 143 SNPs associated with PC1, 91 SNPs associated with PC2, and 149 SNPs associated with PC3 (Supporting Information Table S3). SNPs associated with any of the PCs were widely distributed across all 12 linkage groups in the genome (Figure 2B).

**Figure 2.**
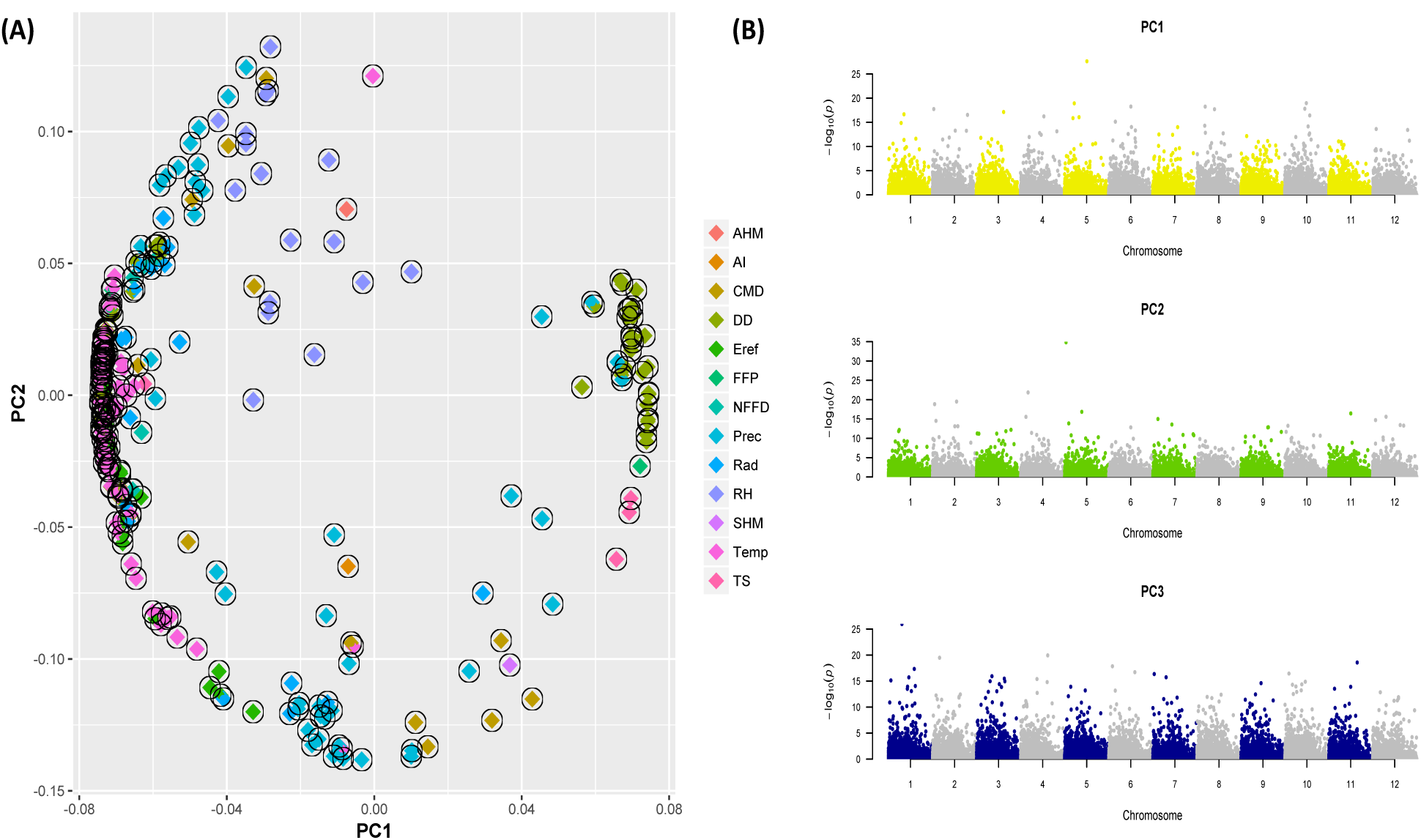
Principal component analysis of environmental variables. Principal components 1 and 2 are shown (Fig.2A). Climate variables full names are shown in Supporting Information Table S1. Figure 2B shows the genomic location of SNPs associated with Principal components of climate variables.

From the 492 individual SNPs showing associations with any climatic variable or Principal Component, 156 SNPs showed clinal patterns with increased or decreased subtle to moderate allele frequency shifts (mean=0.14 ± 0.07, median=0.125) along the longitudinal species’ natural range. Allele frequency of the minor allele increased from east to west in SNPs associated with Climatic Moisture Deficit during winter (CMD_wt), February (CMD02) and March (CMD03), and with Annual Heat Moisture Index (AHM) (Figure 3). SNPs associated with Precipitation as snow (PAS01), Precipitation in September (PPT09), Degree-days above 18 °C, and Number of frost free days during autumn and spring showed an increase in the frequency of the minor allele from west to east. In SNPs associated with PC1 we found 36 SNPs showing increased or decreased allele frequency of the minor allele along longitude; 34 SNPs with PC2; and 59 SNPs with PC3 (Supporting Information Table S4).

**Figure 3.**
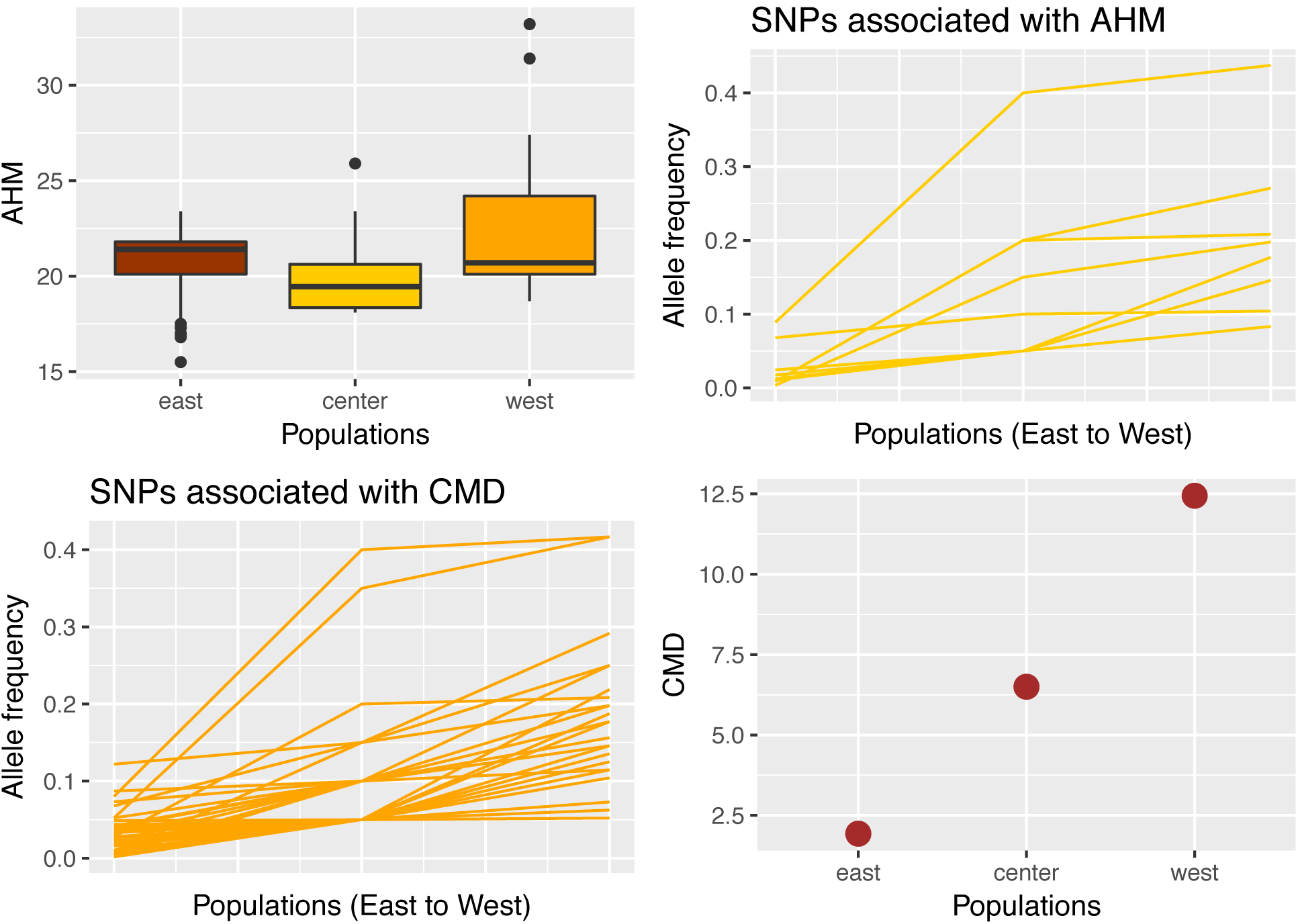
Genetic clines along longitude in which allele frequency of the minor allele increases from east to west in SNPs associated with Annual Heat Moisture Index (AHM) and Climatic Moisture Deficit (CMD).

### Linkage maps

As a result of mapping with ASMap and JoinMap, 17,924 SNPs from the *base* pedigree and 10,995 SNPs from the *qtl* pedigree were mapped in 12 linkage groups in loblolly pine. Pairwise recombination and LOD scores for the *Qtl* and *base* pedigrees’ pseudo-testcrosses can be found in Supporting Information Figure S6. Anchors allowed the construction of average-sex maps using JoinMap. Total lengths were 2158.662 cM for the *base* linkage map and 2141.44 cM for the *qtl* linkage map. The consensus map had a length of 2270.41 cM across 12 linkage groups, and was built with 26,360 SNPs (Table 2, Figure 6, Supporting Information Table S7). These results represent the most complete and dense map ever built for the species (previous map had 3856 markers in Westbrook et al. 2015), and one of the few ultra-dense linkage maps available to date in gymnosperms (others include Norway spruce, Bernhardsson et al., 2019; and *Ginkgo biloba*, Liu et al., 2017). The mapped SNPs were distributed in 18,163 scaffolds in the genome of the species (loblolly pine v2.01 in TreeGenes, treegenesdb.org). Consensus maps had variable measurements of the lowest root mean-squared error (RMSE) standard deviation (mapping conflicts between individuals maps and consensus), ranging from 0.09 to 20.78. Scaffolds’ information of co-located SNPs and assuming all SNPs in the same scaffold were also in the same linkage group, we identified the location of 18,362 more SNPs, resulting in 44,722 SNPs with known positions in linkage groups (17,486 scaffolds) (Supporting Information Table S6). Average number of SNPs among linkage groups was 3726, with the number of SNPs in each linkage group ranged from 3179 to 4148 markers (Supporting Information Table S8). Convergence and map accuracy was evaluated by comparing genome physical location (scaffold ID) and linkage group and by comparing our maps with previously published maps in the species (Martinez-Garcia et al., 2013; Westbrook et al., 2015). Our results indicate that 4% of SNPs had conflicting positions between the physical and linkage positions, suggesting a very small number of SNPs within the same scaffolds were assigned to different linkage groups. These SNPs locations were not considered in further analysis. Comparison between 715 common SNPs in our map and previous maps revealed a convergence of 95.3% of linkage group assignment with Martinez-Garcia et al. 2013 map and 94.2% with Westbrook et al. 2015. Linkage group numbers were chosen to match those at the Martinez-Garcia et al. 2013 linkage map.

**Table 2.** Results of the linkage mapping in loblolly pine showing the length and number of SNPs in each linkage group for individual maps (*base* and *qtl* pedigrees) and consensus map. The ultra­dense consensus map has 26, 021 SNP markers and a length of 2270.21 cM. The root mean-squared error (RMSE) standard deviation gives information about potential mapping conflicts between individuals maps and the consensus map.

| *LG* | *base pedigree* |  | *qtl pedigree* |  | *consensus map* |  |  |
| --- | --- | --- | --- | --- | --- | --- | --- |
|  | *length* | *SNPs* | *length* | *SNPs* | *length* | *SNPs* | *RMSE sd* |
| 1 | 216.386 | 1520 | 180.793 | 844 | 216.39 | 2122 | 20.78 |
| 2 | 205.441 | 1561 | 196.181 | 972 | 198.23 | 2261 | 2.05 |
| 3 | 140.363 | 1538 | 180.977 | 1082 | 187.57 | 2365 | 1.46 |
| 4 | 211.516 | 1465 | 186.414 | 790 | 222.36 | 2022 | 0.09 |
| 5 | 200.858 | 1652 | 177.8 | 955 | 179.19 | 2342 | 1.47 |
| 6 | 196.016 | 1391 | 190.184 | 844 | 196.02 | 2046 | 3.17 |
| 7 | 178.767 | 1278 | 174 | 772 | 178.77 | 1880 | 1.21 |
| 8 | 211.527 | 1628 | 188.5 | 844 | 214.78 | 2252 | 2.4 |
| 9 | 189.513 | 1582 | 156.491 | 844 | 195.6 | 2176 | 0.35 |
| 10 | 136.854 | 1284 | 194.574 | 1065 | 197.53 | 2100 | 15.67 |
| 11 | 119.326 | 1463 | 160.416 | 994 | 131.67 | 2200 | 3.53 |
| 12 | 152.095 | 1562 | 155.107 | 989 | 152.3 | 2255 | 0.73 |
| Total | 2158.662 | 17924 | 2141.437 | 10995 | 2270.41 | 26021 | NA |

### Genomic clusters

We looked for the presence of genomic clusters at four different levels: a) SNPs associated with individual climatic variables; b) SNPs associated with principal components of climate variables; c) SNPs showing high pairwise Fst among populations; and d) SNPs located in the same environmental co-association modules. When evaluating SNPs associated with individual climatic variables, we found that clusters were non-existent (with the exception of a few SNPs associated with Climate Moisture Deficit that co-located at linkage groups 2 and 9; Supporting Information Table S2). In contrast, when evaluating SNPs associated with principal components, we found that larger numbers of co-located SNPs were found to be associated with the same or different groups of environmental variables (PCs) as it was observed in linkage groups 2 (10 SNPs at 190.9 cM), 5 (15 SNPs at 48.76 cM), and 10 (30 SNPs at 19.28 cM). SNPs in LG10 genomic cluster were also included among the top 20% most significant SNPs in the dataset (Figure 4).

**Figure 4.**
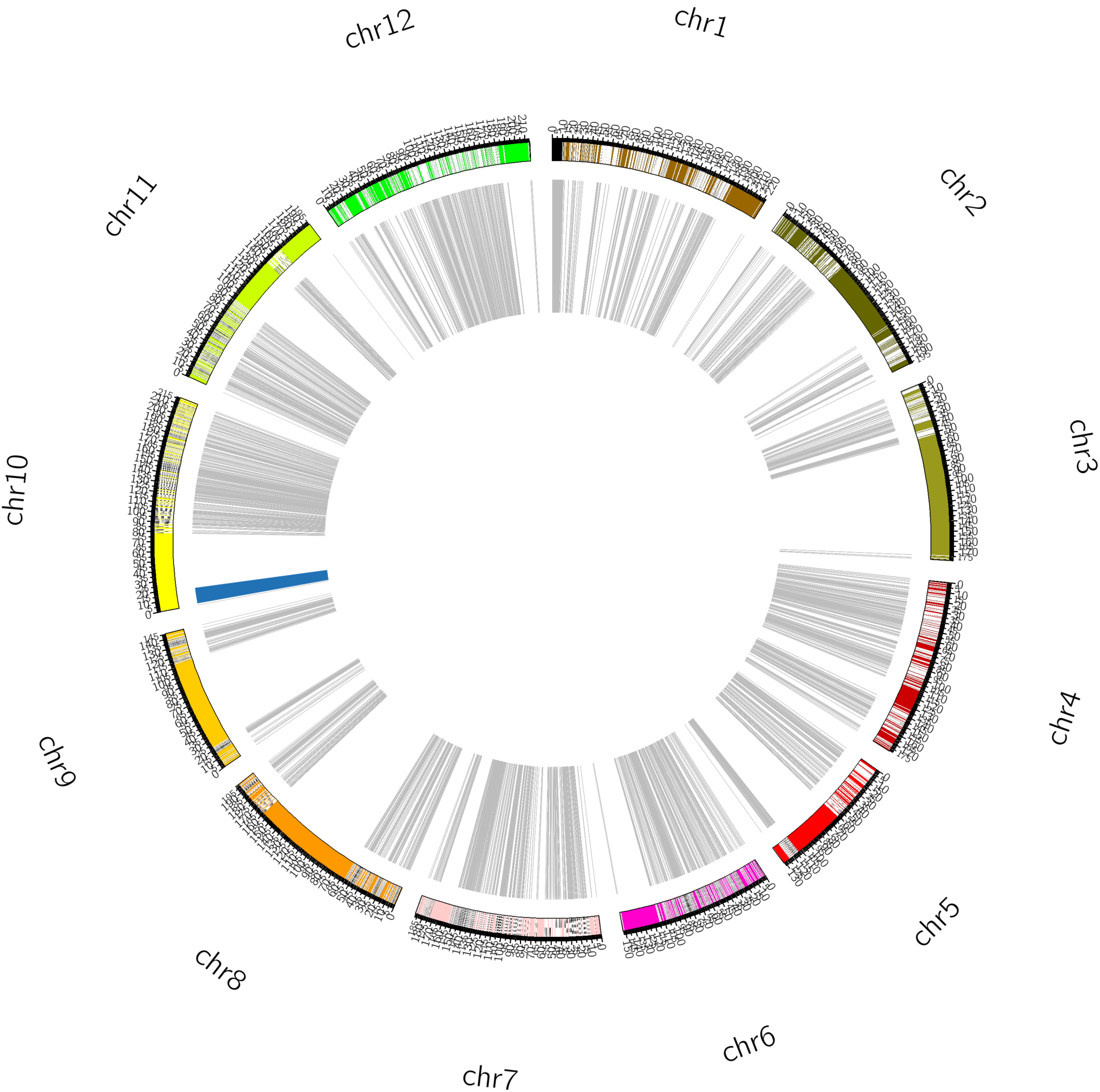
Consensus linkage map containing 26,360 SNPs for loblolly pine. Blue region in the concentric inner circle represent the location of the largest cluster of SNPs associated with principal components of climatic variables. Size of the region was enlarged for easier visualization.

Finally, the evaluation of SNPs showing high population differentiation suggested that many of the SNPs were located in groups with low recombination (co-located or within close proximity) (Supporting Information Table S5). Interestingly, even though many of these SNPs were not found to be associated with any principal component or individual climatic variables, they were located in some of same genomic clusters with SNPs showing associations (Supporting Information Table S5). In the case of SNPs located in the same environmental co-association groups, we found that SNPs located in the aridity module seem to be located in the same scaffold of the species, whereas the SNPs within the humidity module were located in different linkage groups (Supporting Information Figure S4)

## DISCUSSION

Previous studies suggest that population structure of loblolly pine has been mainly shaped by a dual-Pleistocene refugia that separated populations located east and west of the Mississippi river (Wells & Wakeley, 1966; Schmidtling, 2003). In addition to that initial isolation, and in spite of continuous gene flow, east and west populations continued to differentiate as they became adapted to their new distinct environments. Our results suggest that adaptation to climate in the species might have occurred by many changes in the allele frequency of alleles with moderate to small effect sizes, and by the smaller contribution of large effect alleles in genes related to moisture deficit, temperature and precipitation.

### Population structure

Our results suggest the presence of two major genetic clusters (east, west) and a smaller third cluster (center) that extend longitudinally across the species’ natural range (Figure 1A, Supporting Information Figure S1). When K=2, the center and eastern populations are differentiated from the western populations, suggesting the Mississippi river as the major barrier for gene flow. It is being suggested that the Mississippi discontinuity is consistent with a dual­Pleistocene refugial model in which populations in southern Florida and southern Texas later migrated north and expanded their distribution to the current natural distribution of the species (Wells & Wakeley, 1966; Schmidtling, 2003). We found that eastern and western populations present higher pairwise Fst estimates and were therefore more genetically distant than eastern and center, and center and western populations (Figures 1B, 1C and 1D). Despite this differentiation, it is clear that populations were and currently are exchanging gene flow, as general genetic differentiation levels are low suggesting a low population structure. Our results are broadly consistent with previous, smaller scale studies regarding patterns of population structure in loblolly pine (Schimidtling, 2003; Gonzalez-Martinez et al., 2006; Eckert et al., 2010a; Eckert et al., 2010b); and with studies in outcrossing, widespread forest tree species with large population sizes (De La Torre et al., 2014b; Savolainen et al., 2007).

### Water availability is the highest determinant for adaptation of the species

Water stress and temperature variation impose limitations in the survival, growth and productivity of many forest tree species. Loblolly pine is not the exception. Previous physiological and genetic studies have suggested differential responses to temperature and moisture across geographically distant populations of the species (Teskey et al., 1987; Eckert et al., 2010a; Eckert et al., 2010b; Lu et al., 2017). Populations east from the Mississippi river grow faster and taller and are less drought-tolerant than population in the west side (Teskey et al., 1987; Schmidtling, 2001). Although, Lu et al, 2017 found a higher water use efficiency in eastern populations but attributed those results to the young age of trees growing in common gardens, which might be different to mature trees growing in natural populations. It is being suggested that average yearly minimum temperatures of the place of origin affect growth and survival in plantations of the species (Schmidtling, 2001).

In our study, we found that moisture deficit and the relationship between precipitation and temperature seem to be driving the adaptation of the species. Our LFMM results showed that principal component 1, which explained 66.56% of the climatic variation in the dataset, was negatively correlated with Temperature-derived variables, Relative Humidity (RH) and Radiation (Rad); and positively with Degree-days below 0c (DD_0), Degree-days below 18c (DD_18), Precipitation as snow (PAS), Frost-Free period (FFP) and Temperature seasonality variables such as Isothermality (BIO3), Temperature seasonality (BIO4), Annual Temperature Range (BIO7), and Continentality (TD). A total of 143 SNPs were significantly associated with PC1 and many of them showed increased or decreased frequencies of the minor allele along the longitudinal range of the species. Functional annotation of these SNPs included ion transport, fatty acid metabolism, response to auxins, photosynthesis, salt-induced kinases, stress sensing and signal transduction (Supporting Information Table S3). In addition, our GEA results indicate a large number of SNPs associated with Climate Moisture Deficit (CMD_wt, CMD02, CMD03), especially during the winter months; Annual Heat-Moisture Index (AHM) and aridity during the third quarter of the year (AI_Q3); Radiation, Precipitation and Temperature. These SNPs were involved in transport, enzymatic and transcriptional activity (Supporting Information Table S2).

These results are consistent with our previous study in the species that estimate the relationship between expression of xylem development genes and environmental variables. In that study, higher expression levels of MADS box protein, a transcription factor putatively acting as a heat shock protein binding were found with increased levels of climatic moisture deficit in May, and increased radiation in Spring. Also, higher expression levels of Xyloglucan endotransglycosylase 2 (XET-2), an enzyme involved in xylem development (and associated with Laccase 3, Laccase 7 and Phenylalanine ammonia lyase-1), were found with increased radiation during the Spring and decreased precipitation and moisture deficit during the summer (De la Torre et al., 2018).

### Genomic distribution of adaptive alleles

Theory predicts that due to the decrease in fitness with increasing recombination rates, clusters of alleles contributing to adaptive trait variation are frequently located in genomic regions with low recombination (Yeaman, 2013). In our study, however, we didn’t find evidence for genomic clustering or “genomic Islands” in SNPs associated with individual climatic variables, which is consistent with the long-standing view that in conifers linkage disequilibrium decreases rapidly due to their high outcrossing rates leading to high recombination (Neale & Kremer, 2011). Most SNPs showing significant associations with climatic variables or principal components of them had a wide genomic distribution within and among linkage groups, and were present in all the 12 linkage groups of the species. Widespread genomic distributions of adaptive alleles were also found in *Picea mariana* and *Medicago truncatula* (Prunier et al., 2011; Yoder et al., 2014). We only found small haplotype blocks (SNPs closely located with recombination equal or close to zero) in linkage groups 2 and 9 in SNPs associated with Climate Moisture Deficit during winter (CMD_wt) and February (CMD02). Other genomic clustering was not found in SNPs associated with any other climatic variable. However, it is important to mention that although our newly constructed map contains the largest number of SNPs (26k) ever mapped in the species, many of the SNPs showing significant associations with climate locations could not be located within linkage groups. An even higher density linkage map or a chromosome-scale reference genome would be required to confidently locate most or all of the SNPs associated with climatic variables.

Interestingly, when evaluating SNPs associated with Principal Components and environmental co­association modules, we found larger numbers of co-located SNPs. This suggests that adaptation to climate in loblolly pine may have occur as a complex process in which different selection pressures are more likely to act on groups of genes associated with an aspect of climate rather than on individual climatic variables. These “co-adapted” complexes of genes may buffer against gene flow coming from maladaptive alleles from geographically proximal but climatically different locations, maintaining polymorphisms across the species’ natural range (Holliday et al., 2016). Increased clustering of outlier loci was found across altitudinal gradients with high gene flow between populations of *Populus trichocarpa*, suggesting adaptation with gene flow might have occurred by divergence hitchhiking of physically proximate alleles in the species (Holliday et al., 2016). Evidence for recurrent hitchhiking was also found in *Capsella grandiflora*, an outcrossing species with large effective population size and low levels of linkage disequilibrium (Williamson et al., 2014).

In addition to SNPs adapting to the same aspect of climate or PC, we also found a smaller group of co-located SNPs associated with different groups of environmental variables (PCs) as it was observed in linkage groups 2, 5 and 10 (Figure 6). In addition, a large cluster of SNPs was found at position 19.28 cM in linkage group 10, where 30 different SNPs associated with PC1, PC2 and PC3 were co-located. It is important to mention, however, that map accuracy of linkage group 10 might have been lowered by some discrepancies between individuals maps and the consensus map, as it was suggested by a higher RMSE (root mean-squared error) than in other linkage groups. Our results suggest that SNPs located in the aridity co-association module seem to be located in the same scaffold of the species, whereas the SNPs within the humidity module were located in different linkage groups. Physical linkage among loci adapting to different aspects of climate was also found in *Pinus contorta*, while studying modules of co-associated SNPs (Lotterhos et al., 2018). Both Lotterhos et al. 2018 and this study suggest a complex genomic architecture of local adaptation in conifer species, in which the extent of physical linkage among loci is just one of the factors contributing to the species’ evolutionary response to changes in the climate.

Perhaps our most interesting finding in relation to the genomic architecture in loblolly pine was that in spite of the rapid decay of LD, we found genomic regions of low recombination in three of the twelve linkage groups. These genomic regions showed a variety of SNPs associated with either PC1, PC2, PC3, or Climate Moisture Deficit and showed high population differentiation. Increased linkage disequilibrium between these climate-associated alleles of small effect may have prevented their swamping by gene flow and may have promoted their contribution to adaptive trait divergence (Yeaman & Whitlock, 2011; Yeaman, 2015). In the case of the SNPs located in the genomic cluster in linkage group 5 (48.04-48.84 cM), we found a correlation between increased population differentiation and decreased diversity (R^2^= 0.3, P<0.05). A negative relationship between recombination rate and genetic differentiation is a common signature of linked selection and has been observed in several plant species including *Populus tremula* and *P. tremuloides* (Slotte et al., 2014; Wang et al., 2016). In addition, two of these genomic clusters (groups 2 at 189.62-191.79 cM and 5 at 48.04-48.84 cM) were previously considered as “metabolic hotspots” because they harbor SNPs associated with several metabolites such as pelargonic acid, threonine, and other metabolites of unknown origin (De La Torre et al., 2018).

### Adaptive alleles- globally occurring or localized?

An important question in population genetics is whether alleles conferring adaptation are globally occurring alleles or localized ones. If natural selection favors specific alleles in specific locations, it is expected that these would be common in these geographic locations but rare in others. On the contrary, if natural selection removes alleles that are deleterious in one location but neutral in others, we would expect to find high-frequency alleles across the species’ range (Fournier-Level et al., 2011). Localized or private alleles with narrow geographic distribution in which one allele has rapidly driven to high frequency in the population may be a result of a hard-selective sweep, as it is being observed in natural populations of *Arabidopsis thaliana* (Hancock et al., 2011). Alternate alleles might be favored in different environments leading to antagonistic pleiotropy that can result in local adaptation and the maintenance of genetic polymorphisms by natural selection. In a different scenario (conditional neutrality), alleles may be under positive selection in one environment but neutral in others (Anderson et al., 2013).

Our study found that putatively adaptive alleles in loblolly pine were widely distributed across the species’ natural range rather than localized ones. In fact, the presence of private alleles (only present in one population) was not observed in any of the SNPs showing associations with climate variables. Globally occurring alleles had varying frequencies in which the frequency of the minor allele increased from east to west in SNPs associated with Climatic Moisture Deficit during winter (CMD_wt), February (CMD02) and March (CMD03), and with Annual Heat Moisture Index (AHM); whereas SNPs associated with Precipitation as snow in January (PAS01), Degree-days above 18 °C during winter, and Number of frost free days during autumn and spring, showed an increase in the frequency of the minor allele from west to east. In *Arabidopsis thaliana* populations, SNPs associated with relative humidity also had widespread distributions (Hancock et al., 2011). With the absence of fitness measurements, we cannot tell if loblolly pine individuals carrying these alleles are fitter in one environment or the other. However, the fact that the direction of the increase of the minor allele is coincident with increasing levels of the associated climatic variable suggests that these individuals may be locally adapted in that environment.

### Allele frequency shifts at many adaptive loci

Local adaptation in natural populations may arise by differential selection pressures across heterogeneous environments in which the targets of selection may change from one environment to another. As a consequence of this, different combinations of alleles might be favored in different environments and maintained as stable polymorphisms, or experience “partial” or “soft” sweeps due to selection acting on standing variation (Hermisson & Pennings, 2005; Yoder et al., 2014). Recent studies have found a largely polygenic basis of adaptation in natural populations, in which trait variation is controlled by many loci of small effect and adaptation is characterized by subtle to moderate shifts in allele frequencies (Pritchard et al., 2010a, 2010b; Berg & Coop, 2014). With strong diversifying selection and high gene flow, considerable trait divergence may evolve with small allele frequency changes at individual loci (low Fst) but high between-population covariance in allele effect sizes (Latta, 1998; Le Corre & Kremer, 2003, 2012). Adaptation to climate variation via selection on polygenic traits and/or small allele frequency shifts has been observed in *Medicago truncatula*, *Picea glauca*, *Fagus sylvatica* and *Maccullochella peelii* (Csillery et al., 2014; Yoder et al., 2014; Hornoy et al., 2015; Harrison et al., 2017). In *Picea glauca*, small to moderate shifts in the allele frequency of putatively climate-adaptive genes was found in response to recent selection and high gene flow among populations (Hornoy et al., 2015).

In our study, we found a large number of SNPs with small to moderate effect sizes associated with climatic variables or combinations of them. In most of these SNPs, we found subtle to moderate shifts in allele frequencies across different environments, in which in many cases the increase in the frequency of the minor allele mirrored an increase of the climate variable along longitudinal gradients. These SNPs were largely captured by our GEA study, which, in contrast to Fst outlier tests has been suggested to detect signals associated with smaller allele frequency shifts (Hancock et al., 2010). Coincidently, our Fst outlier test failed to detect the signal of selection in these weakly selected loci involved in local adaptation of loblolly pine. In polygenic traits, most loci involved in local adaptation will experience weak selection, therefore they will not be substantially more differentiated than expected of neutral loci (Whitlock & Lotterhos, 2015; Le Corre & Kremer, 2012). Loci highly associated with expression and disease resistance and previously suggested to be under balancing selection (De La Torre et al., 2018) were also not identified by the Fst outlier analysis, probably because the OutFLANK procedure is not accurate in the left tail of the Fst distribution (Whitlock & Lotterhos, 2015). The little difference in Fst across associated versus randomly chosen SNPs may also suggest that natural selection is not driving large-scale adaptive differences among lineages of loblolly pine. Instead, genotypes are being favored by natural selection across different environments regardless of their ancestry (Eckert et al., 2010b).

Responses to selection that arise from standing genetic variation rather than new mutations or that are relatively recent for fixation to have occurred leave a fainter molecular signature (Hermisson & Pennings, 2005; Hohenlohe et al., 2010). These partial, soft sweeps often leave a signature of reduced haplotype or nucleotide diversity and extended linkage, as we found in our LG 5. Considering the low mutation rates in conifers (De La Torre et al., 2017), it is likely that many of these changes in allele frequencies might have been facilitated by the great levels of standing genetic variation rather than by de novo mutations in loblolly pine. Simulations studies have suggested that good levels of standing genetic variation are required when local adaptation occurs by alleles of small effect (Yeaman, 2015). In addition, the long generation times in conifers and relatively recent migration from refugia of the species, might have contributed to the genetic architecture we see today, in which most beneficial alleles are still segregating in the populations and have not reached fixation. We therefore conclude that local adaptation to climate in loblolly pine might have occurred by many changes in the allele frequency of alleles with moderate to small effect sizes, and by the smaller contribution of large effect alleles in genes related to moisture deficit, temperature and precipitation.

## ACKNOWLEDGEMENTS

This project was supported by the U.S. Department of Agriculture/National Institute of Food and Agriculture (“PineRefSeq”; 2011-67009-30030) awarded to D.B.N at the University of California, Davis. The authors would like to thank Chuck Burdine, Patrick Cumbie and Dana Nelson for sample collection; and members of the Neale’s Lab Annarita Marrano, Sara Montanari and Pedro Martinez for suggestions on linkage mapping.

## DATA ACCESIBILITY

Consensus linkage maps and genotyping data will be available in public database repositories such as TreeGenes and Dryad after the acceptance of the paper.

## AUTHOR CONTRIBUTIONS

D.N and A.D.L.T designed the research study, A.D.L.T did all the data analyses and wrote the manuscript, D.N reviewed and commented the final version of the manuscript.

## SUPPORTING INFORMATION

**Table S1**. List of climatic variables showing significant associations with one or more environmental variables tested in this study.

**Table S2**. Results of the GEA analysis based on 87k genome-wide SNPs using GAPIT R package. Variables include: climate variable, SNP marker, physical location (scaffold id), physical position (scaffold position), consensus linkage group (LG) and position (cM), consensus_physical_linkage (linkage group is known but not position), maf (minor allele frequency), R^2^ (percent of variation explained), p-value (p-value before correction for multiple testing), adjusted_p-values (p-value after correction for multiple testing), annotation and annotation id, functional annotation, and species (reference for functional annotation).

**Table S3**. Results of the GEA analysis using the program LFMM. Only associated SNPs within the top 20% with the highest −log_10_(p-value) are shown. The first three principal components of environmental data were tested for associations with 87k genome-wide SNPs. Variables include: climate variable, SNP marker, physical location (scaffold id), consensus linkage group (LG) and position (cM), consensus_physical_linkage (linkage group is known but not position), maf (minor allele frequency), R^2^ (percent of variation explained), p-value (p-value before correction for multiple testing), adjusted_p-values (p-value after correction for multiple testing), annotation and annotation id, functional annotation, and species (reference for functional annotation).

**Table S4**. Genetic clines along longitude for SNPs showing significant associations with environmental variables or principal components of them. Variables include: climate variable, SNP, allele frequencies in the east, center and west populations, allele frequency differential, direction of increase in allele frequency, maf (minor allele frequency), p-value (p-value before correction for multiple testing), adjusted_p-values (p-value after correction for multiple testing), annotation and annotation id, functional annotation, species (reference for functional annotation),

**Table S5**. Genomic clusters of SNPs showing significant pairwise population differentiation. Variables included are: SNP, physical location (scaffold id), physical position (scaffold position), consensus linkage group (LG) and position (cM), Pairwise Fst between pairs of populations, nucleotide diversity for each population, annotation and annotation id.

**Table S6**. Genomic location for all the 87k SNPs in this study taken from our newly constructed ultra-dense linkage maps in loblolly pine. Variables indicate: physical location in the genome (scaffold id), consensus linkage group (LG) and position (cM), and consensus_physical_linkage (linkage group is known but not position),

**Table S7**. Number of SNPs identified in each linkage group after construction of the consensus ultra-dense linkage map and incorporation of physical (scaffold) location.

**Figure S1**. Principal component analysis of genotyping data for 87k SNPs suggest the presence of three major genetic clusters (from left to right: east, center and west) that extend longitudinally across the natural distribution of loblolly pine.

**Figure S2**. FastStructure cluster plots from K=2 to K=5 show two major genetic clusters (east and center in blue, and west in orange) in the natural distribution of loblolly pine.

**Figure S3**. Significant correlations between individual environmental variables and Principal Component 1 (PC1).

**Figure S4**. Results of environmental co-association network analysis. Hierarchical clustering of top-significant (R2>0.4) associations between SNP allele frequencies and environmental variables shows two main clusters, right cluster: related to aridity (temperature variables, radiation and degree-days above 18°C) and a left cluster: mainly associated with humidity and cold (precipitation during winter).

**Figure S5**. Heatmap showing correlations among individual environmental variables. Large cluster in blue (top right) is composed by associated variables NFFD, Radiation (Rad and MAR), Temperature-related variables, Degree-days above 18°C during the summer, Evaporation (Eref), and Degree-days above 5°C.

**Figure S6**. Pairwise recombination and Linkage disequilibrium in loblolly pine; (A) recombination (left side) and LOD scores (right side) in *Qtl* pedigree pseudo-testcross 1; (B) recombination (left side) and LOD scores (right side) in *Qtl* pedigree pseudo-testcross 2; (C) Map recombination fraction, map distance, LOD scores, estimated recombination fraction and number of recombination events in each linkage group in the *Qtl* pedigree pseudo-testcross 1.

